# Maternal control of spontaneous dormancy termination in *Daphnia pulex*

**DOI:** 10.1101/2022.12.13.519803

**Authors:** Robert J. Porter, Grace M. Gutierrez, Karen B. Barnard-Kubow, Alan O. Bergland

## Abstract

This study examined maternal influence and life-history consequences of diapause termination timing in *Daphnia pulex*. We raised clonal isolates of *D. pulex* in mesocosms and observed hatching rates prior to and after exposing embryos to a cold shock. A substantial proportion of individuals hatched early, prior to the cold shock. We found that siblings from the same ephippium were more likely than expected by chance to emerge at the same time, even after dissection and separation, suggesting the presence of a maternal effect that influences diapause duration. We also found that for individuals who emerged early, the time to first reproduction was significantly delayed, and individuals produced fewer resting embryos in subsequent generations. We suggest that early diapause termination may be driven by maternal effects to generate offspring that emerge from dormancy at different times.

## Introduction

Dormancy, the brief cessation of development, is critical for the survival of organisms that live in harsh and ever-changing environments. One dormancy strategy is diapause, a period of arrested development in times of unfavorable environmental conditions (Hand et al., 2016; Hairston & Kearns, 2002; Lopes et al., 2004). Some organisms, such as killifish (Dolfi et al., 2019) and rotifers (Ricci, 2001) produce diapausing resting embryos capable of surviving desiccation and freezing. Typically, resting embryos resume development and emerge when environmental conditions return to being hospitable, at the beginning of the next growing season. However, variation in dormancy strategy can occur in response to heterogeneity in habitat quality over time (ten Brink et al., 2020). In environments where habitat quality predictably and drastically declines and resumes, the timing of dormancy is strictly reinforced by selection (Wagner & Simons, 2009). When habitat quality does not change drastically or varies unpredictably, the strict timing of dormancy is no longer enforced by selection, and consequently individuals can terminate dormancy early (Satake et al., 2001) or not enter dormancy at all (Ensslin et al., 2018). Variation in the timing of dormancy can cause individuals to emerge at different times, and experience different environments (Postma & Ågren, 2022). We hypothesize that early termination of dormancy plays a major role in life history differences between individuals who emerge at different times.

The cladoceran *Daphnia* is a prime example of an organism found in many changing environments, from permanent lakes subjected to near freezing temperatures (Grosbois et al., 2017), to fully ephemeral ponds that frequently become inhospitable (Lynch, 1983). Under favorable environmental conditions, *Daphnia* reproduce parthenogenetically. This form of asexual reproduction results in nearly identical clonal female offspring (Hebert & Ward, 1972, Flynn et al., 2017). However, in order to survive unfavorable environmental conditions such as stress caused by freezing or drought, *Daphnia* produce resting embryos that are deposited within an ephippium--a protective, drought-resistant envelope (Hiruta & Tochinai, 2014). Ephippia hold up to two resting embryos but may contain one or no embryos depending on mating success (Winsor & Innes, 2002). Ephippia are produced in place of an asexual clutch, which would typically contain 2-40 offspring (Gliwicz & Lampert, 1994). Therefore, to produce an ephippium, *Daphnia* sacrifice immediate population growth to invest in future sexual offspring (Gerber et al., 2018). This strategy, while costly in the short term, ultimately allows *Daphnia* populations to propagate following the cessation of harsh conditions, as well as generate recombinant offspring. Resting embryos within the ephippium can survive food scarcity, hypoxia, drought, and freezing temperatures until conditions for hatching become favorable again (Radzikowski et al., 2018). As such, it is not expected for resting embryos to emerge prior to experiencing harsh conditions.

The specific conditions that induce *Daphnia* to emerge from their diapausing state have been well described. Laboratory experiments have found that stimuli such as photoperiod (Pancella & Stross, 1963), temperature (Vandekerkhove et al. 2005), and ephippia decapsulation (Davison, 1969, Paes et al., 2016) affect the rate of diapause termination. Numerous studies have found that incubation of the resting eggs in low temperature and dark conditions, followed by exposure to warm temperatures and bright light, increases diapause termination of ephippial offspring (Davison, 1969; Schwartz & Hebert, 1987; Vandekerkhove et al., 2005; Shabani et al., 2012; Luu et al., 2020). However, few studies have investigated the early termination of diapause prior to experiencing these conditions in *Daphnia* (Cáceres, 1998), nor the life history effects associated with early diapause termination.

To investigate the phenomenon of early diapause termination in *Daphnia*, we observed diapause termination rates from two clonal isolates of *D. pulex*. This study investigated (i) the role of maternal effects on embryonic diapause, (ii) the impact of early diapause termination on offspring life history traits, and (iii) the role of dormancy strategy on transgenerational reproductive effects. From this study, we demonstrate the existence of early diapause termination in *D. pulex* and suggest that maternal effects are a major driving force in the timing of dormancy.

## Materials and Methods

### Study Population

*D. pulex* were collected from a semi-permanent pond in Dorset, England in the Kilwood Nature Reserve (50.64, - 2.09) in the spring of 2017. Pond ephemerality was determined using trail cameras, and multi-year survey of the genetic composition of *D. pulex* (Barnard-Kubow et al., 2022). Live *Daphnia* samples were transported back to the University of Virginia, where clonal lineages were established from wild female isolates. Lineages were maintained in 18C and 16:8 light:dark conditions and raised in 250 ml glass jars where they underwent asexual reproduction. The jars were maintained in hard artificial pond water (ASTM) (Standard, 2007) and fed with lab-grown *Chlorella vulgaris* obtained from UTEX (strain #4223).

### Mesocosm Conditions

15 liter mesocosms filled with ASTM were established in the spring of 2019, and inoculated with one of either two *D. pulex* lineages (D8515, D8222), which were established from two separate field isolates of the same wild superclone lineage (superclone C, Barnard-Kubow et al., 2022). Clonal isolates D8515 and D8222 are nearly genetically identical, although there are a limited number of genetic differences that have arisen due to mutation accumulation and gene conversion (Barnard-Kubow et al., 2022). The mesocosms were seeded with ∼60 adult females, and were placed in a Percival incubator (Model Number: 1166NL) at 18C 18:6 light:dark conditions using LED lights (∼1500 lux, Royal Pacific 4311WH). Mesocosms were fed with 95,000 cells/mL *C. vulgaris* every Monday, Wednesday, and Friday for the first five weeks, then 142,000 cells/mL until week eight. Mesocosms were tracked for eight weeks and sampled weekly to determine age and reproductive demographics.

### Ephippial Hatching

Ephippia were collected on a weekly basis from mesocosms of both clonal lineages and stored individually in clear 96 well plates filled with ASTM at 18C under long day conditions (16L:8D) for four weeks (N=2615). A subset of ephippia (N=763) were labeled according to whether they were found to be floating on the surface, or fully submerged. Immediately after collection, a portion of ephippia (N=402/2615, 15.37%) were manually decapsulated using fine metal needles in order to provide a general estimate of the number of eggs (Paes et al., 2016). The decapsulated embryos obtained from the dissected ephippia were also placed under the same hatching conditions as the non-dissected ephippia. Each plate was checked every two to three days under lower light using a Leica S8 APO microscope to determine hatch rate.

As individuals began to develop, they were removed from the plates and isolated in 50ml jars with ASTM to establish new isofemale lines. Any individuals that had not hatched after four weeks were incubated for four more weeks in a cold treatment in complete darkness at 4°C. After the four weeks in 4°C, individuals were then removed and placed once more under hatching conditions (18C, 18L:8D) to develop for a further two weeks, at which point no more individuals developed. All undissected ephippia were dissected at the end of the experiment, and embryos counted to determine the total number of embryos produced over the course of the experiment (N=1931 embryos).

### Jar Censusing

We followed 1561 sexually produced ephippial hatchlings over the course of 11 weeks. Within this number, a total of 233 sibling pairs were followed to determine maternal effects, including 60 sibling pairs from ephippia that were dissected. All sexually produced individuals that developed were isolated in 50ml jars with ASTM and were checked weekly for five weeks to determine survival and reproduction. After reproduction was established, individuals were moved to 250ml jars, and were then further checked at 5, 8, and 11 weeks for male and ephippia production. The presence of males was established by random sampling of 9-12 sub-adult individuals. Any individuals that failed to reproduce by 5 weeks were marked as sterile. Any ephippia produced by within isofemale lines were collected and dissected to establish the number of embryos produced, as the presence of filled ephippia could be used as a metric of the presence of males.

### Statistical Analysis

Statistical analyses were performed using R version 3.6.2 (R Core Team, 2019). We performed analysis of deviance to test for the factors that affect early hatching and to test whether early hatching affects other traits in the offspring. When the dependent variable was binary or a proportion (survival, hatch rate), we used a generalized linear model with binomial error structure (Models 1, 2, 3, 5, 6, Table 1). When the dependent variable was count based (number of ephippia produced), we used a Poisson error (Model 4, Table 1).

**Table 1:** Models and ANOVA Results.

| (Model 1) Proportion early hatching ~ Clone * Mesocosm Age |  |  |  |  |  |
| --- | --- | --- | --- | --- | --- |
|  | DF | Deviance | Resid. DF | Resid. Dev | Pr(>Chi) |
| Clone | 1 | 26.62 | 8 | 68.183 | 2.486E-07 |
| Mesocosm Age | 1 | 34.03 | 7 | 34.156 | 5.434E-09 |
| Clone:Mesocosm Age | 1 | 1.595 | 6 | 32.561 | 0.2066 |
| (Model 2) Survival to reproduction ~ Hatching Time * Dissection Status + Clone |  |  |  |  |  |
|  | DF | Deviance | Resid. DF | Resid. Dev | Pr(>Chi) |
| Hatching Time | 1 | 1.583 | 6 | 9.0374 | 0.2084 |
| Dissection Status | 1 | 0.1002 | 5 | 8.9372 | 0.7515 |
| Clone | 1 | 3.647 | 4 | 5.2906 | 0.05619 |
| Hatching Time:Dissection | 1 | 4.706 | 3 | 0.5843 | 0.03005 |
| (Model 3) Proportion Reproduced ~ Hatching Time * Jar Age + Clone |  |  |  |  |  |
|  | DF | Deviance | Resid. DF | Resid. Dev | Pr(>Chi) |
| Hatching Time | 1 | 4.14 | 14 | 950.26 | 0.04199 |
| Jar Age | 1 | 567.1 | 13 | 383.13 | < 2.2e-16 |
| Clone | 1 | 20.8 | 12 | 362.33 | 5.092E-06 |
| Hatching Time:Week | 1 | 5.02 | 11 | 357.3 | 0.02499 |
| (Model 4) Ephippia production ~ Hatching Time * Clone |  |  |  |  |  |
|  | DF | Deviance | Resid. DF | Resid. Dev | Pr(>Chi) |
| Hatching Time | 1 | 265.6 | 644 | 3419.9 | < 2.2e-16 |
| Clone | 1 | 17.78 | 643 | 3402.1 | 2.478E-05 |
| Hatching Time:Clone | 1 | 4.977 | 642 | 3397.1 | 0.02569 |

(Model 5) Ehippia fill rate ~ Hatching Time \* Clone
|  | DF | Deviance | Resid. DF | Resid. Dev | Pr(>Chi) |
| --- | --- | --- | --- | --- | --- |
| Hatching Time | 1 | 0.0622 | 318 | 2745.1 | 0.8031 |
| Clone | 1 | 0.7499 | 317 | 2744.4 | 0.3865 |
| Hatching Time:Clone | 1 | 16.79 | 316 | 2727.6 | 4.166E-05 |

(Model 6) Early hatching ~ Buoyancy \* Clone
|  | DF | Deviance | Resid. DF | Resid. Dev | Pr(>Chi) |
| --- | --- | --- | --- | --- | --- |
| Buoyancy | 1 | 8.556 | 2 | 38.243 | 0.003445 |
| Clone | 1 | 26.88 | 1 | 11.362 | 2.164e-07 |
| Buoyancy:Clone | 1 | 11.36 | 0 | 0 | 0.0007496 |

## Results

This study investigated the rate of early diapause termination in lab-reared *D. pulex*. We characterized the role of maternal effects on the rate of early diapause termination, as well as the effects of early diapause termination on offspring’s life history traits. We found that many hatchlings (N=413/1561, 26.46%) emerged from diapause without any obvious changes in environmental cues.

### Maternal effects on early hatch rate

To assess whether maternal effects influenced the timing of hatching, we examined 233 sibling pairs that were deposited into the same ephippia. These individuals are presumed to be full-siblings (Hiruta & Tochinai, 2014). Because all of the ephippial mothers are from the same clonal lineage, embryos produced by different mothers are all effectively full-siblings. If there is maternal control of diapause variation, we would expect embryos from the same ephippia – and thus same individual mother – would be more likely to have the same hatching behavior than expected based on the average rate of early diapause termination across all embryos. We found that embryos from the same ephippium were more likely to emerge at a similar time: there were fewer embryos from the same ephippium where one hatched prior to cold treatment and one hatched after cold treatment than would be expected by chance (df= 1, *χ*^2^ = 122.1, p=2.195e-28, Table 2). Moreover, the hatching dynamics of embryos from the same ephippia were similar regardless of whether we dissected ephippia (N= 60 pairs) or let embryos naturally hatch (N=173 pairs), ruling out the possibility that the development of one embryo triggered the development of the other (Fig 1A). We assessed whether the maternal environment affected the early hatching rate as a function of mesocosm age. Over the 8 sample weeks, the density of individuals increased from 60 individuals to several thousand (Barnard-Kubow et al., 2022). To assess the early hatching rate of the two clonal lineages, we collected ephippia weekly from the mesocosms, and found that the rate of early hatching increased as mesocosms became older (df= 1, *χ*^2^ =34.03 p=5.434e-09, Fig 1B, Model 1).

**Table 2:**
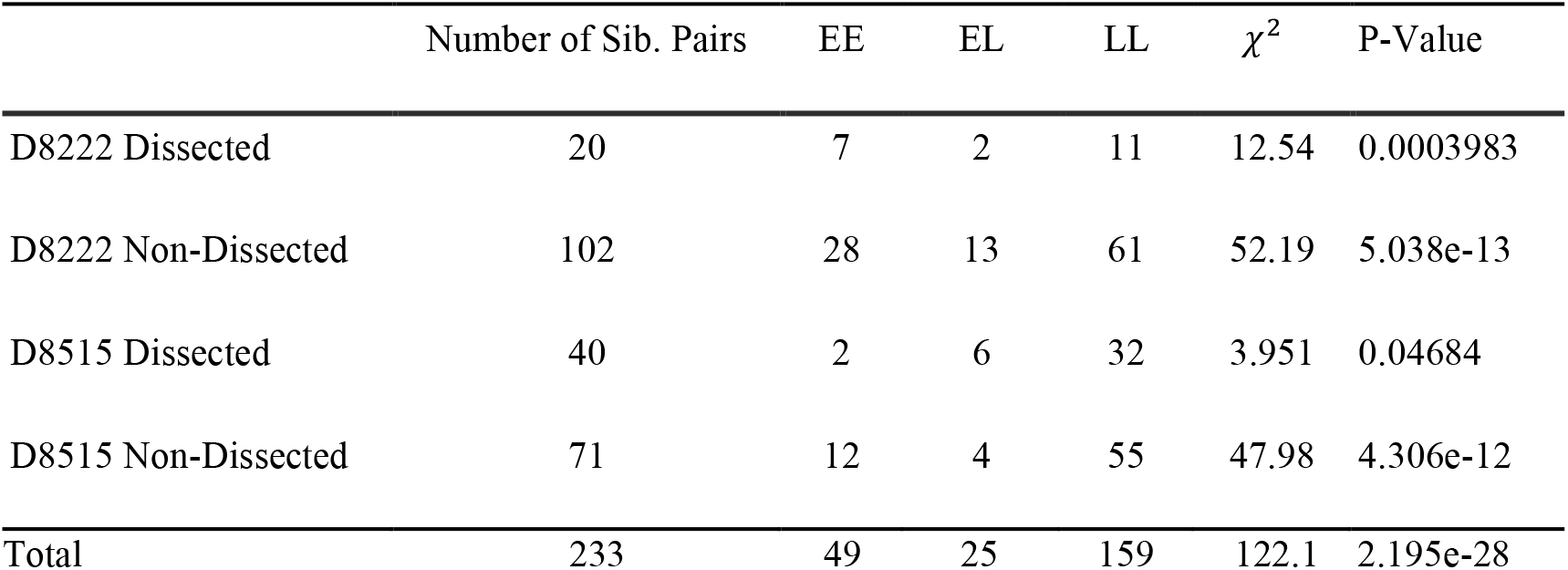
Individuals tend to hatch out with their immediate sibling in similar conditions (EE, LL) than expected by chance. There were fewer sibling pairs where one individual hatched early, and one had hatched late (EL) than expected by chance.

**Figure 1.**
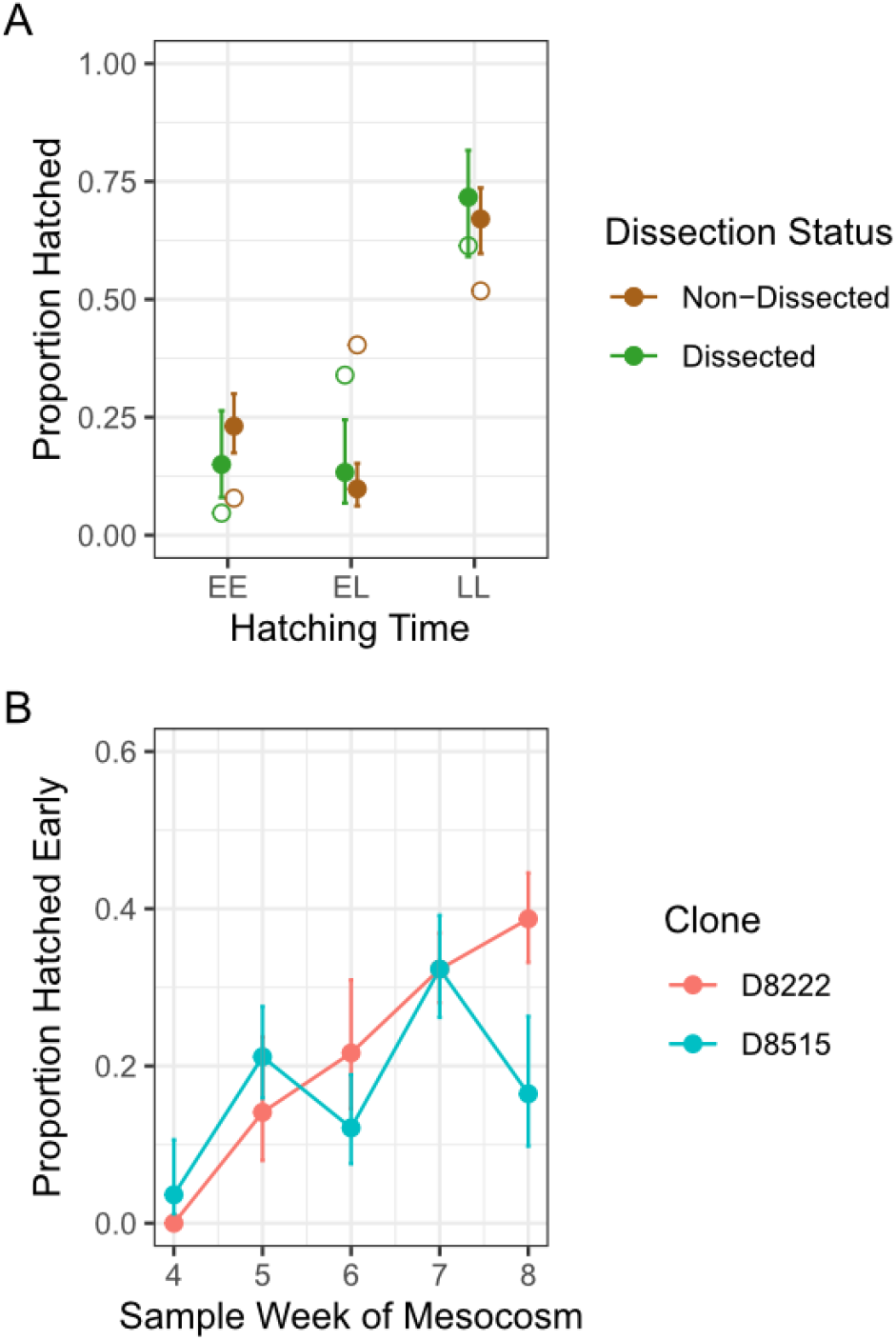
Maternal environment and maternal cues affect early hatching. (A) Proportion of sibling pairs that have hatched out based on treatment. EE - both siblings emerged early; EL - One sibling emerged early, the other late; LL - Both siblings emerged late. Individuals tend to hatch out with their immediate sibling. Error bars represent 95% confidence intervals calculated using standard error. D8222 week 4 had too few individuals to calculate statistics (N = 0/2 hatched early). Open circles represent expected values based on the marginal rate of early hatching; filled circles represent the observed values (B) Proportion of individuals that hatched early over the course of the 8 week mesocosm. The rate of early diapause termination increased as the mesocosms aged (df= 1, *χ*^2^ = 34.03, p=5.434e-09, Model 1). Each line represents a clonal isolate, with error bars representing 95% confidence intervals calculated using standard error.

### Hatchling survival and reproduction

Early hatching did not significantly affect individual survival to reproduction (df=1, *χ*^2^ =1.583, p= 0.2084, Fig 2A, Model 2). Moreover, whether individuals were dissected or naturally hatched did not significantly affect survival (df=1, *χ*^2^=0.1002, p=0.7515, Fig 2A, Model 2). Individuals that hatched early took a slightly, but significantly, longer amount of time to reach reproduction compared to those that hatched late (df=1, *χ*^2^ =4.14, p=0.04199, fig. 2B, Model 3).

**Figure 2.**
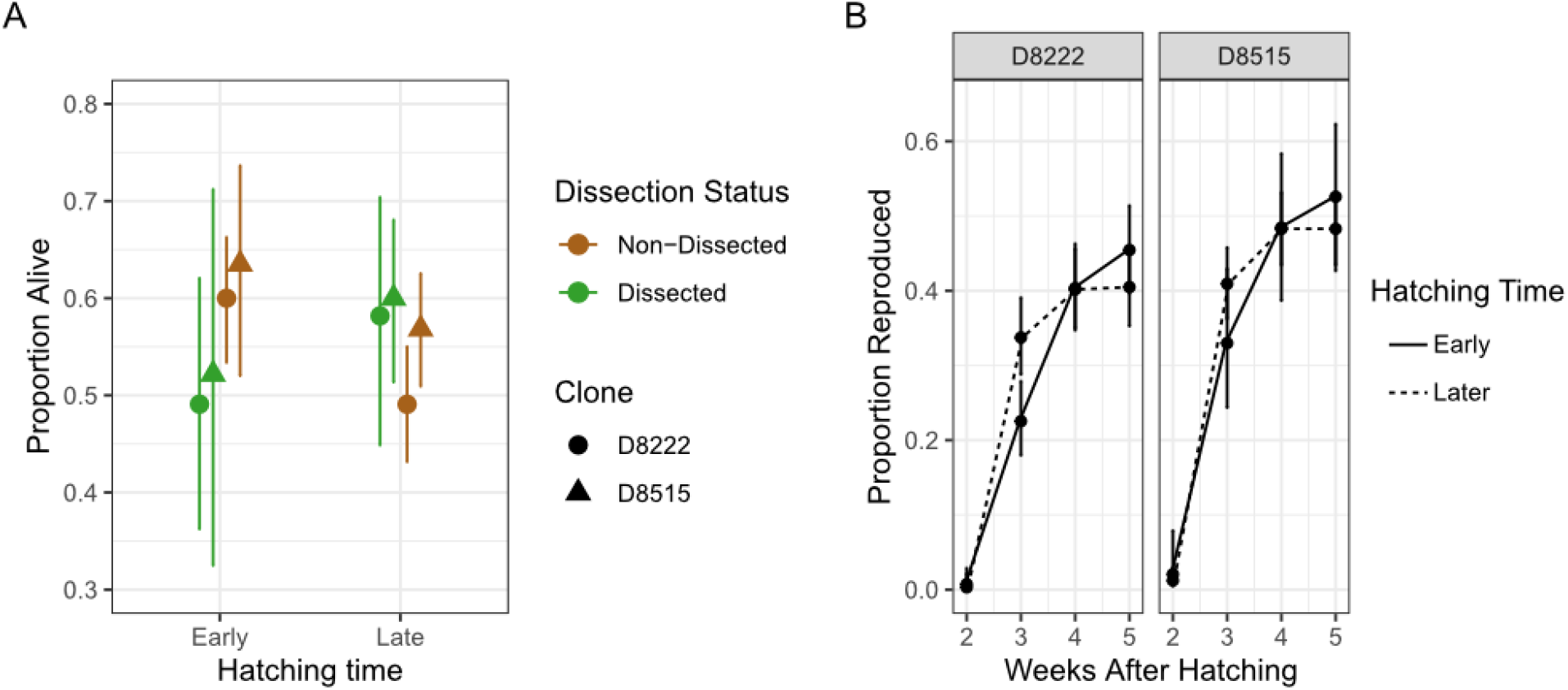
Effects of diapause duration on life history traits. **(A)** There were no major differences in survival over the first five weeks between individuals that hatched out early or late (df=1, *χ*^2^ =1.583, p= 0.2084, Model 2), nor did dissection affect survival (df=1, *χ*^2^=0.1002, p=0.7515, Model 2). (B) Time to reproduction differed significantly between treatments. The time to first reproduction was slightly, but significantly slower in individuals that had hatched early (df=1, *χ*^2^ =4.14, p=0.04199, Model 3).

### Transgenerational effects of early hatching

To determine transgenerational effects, we measured the number of ephippia produced by hatchlings and their immediate offspring and assessed the ephippial fill rate as a proxy for male production and mating success. Lineages produced by offspring that emerged early produced significantly fewer ephippia than individuals that hatched out late (df=1, *χ*^2^ =265.6, p=2.2e-16, Fig 3A, Model 4). The two parental clonal isolates also differed in ephippia production (df=1, *χ*^2^ =17.78, p=2.478e-05, Model 4). Additionally, we found a slight difference in the number of ephippia produced between each clonal isolate, depending on whether the replicate hatched out early or late (df=1, *χ*^2^ =4.977, p=0.02569).

**Figure 3:**
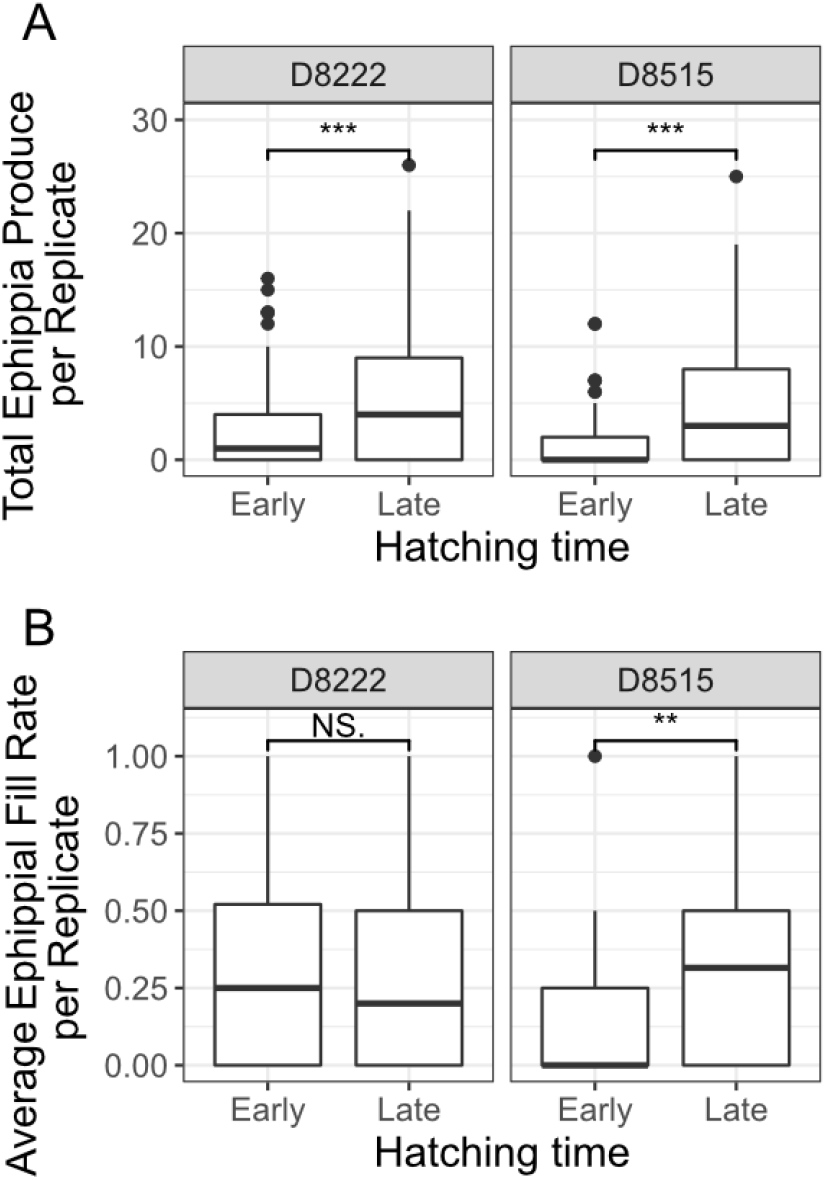
Early hatching affects some offspring life-history traits. (A) Average ephippia per jar significantly increased in post-treatment individuals (df=1, *χ*^2^ =265.58, p=2.2e-16, Model 4). (B) Average ephippia fill rate did not change significantly between treatments in D8222 but did in D8515.

We did not detect any significant effect of hatching time on ephippial fill rate (df=1, *χ*^2^ =0.06227, p=0.8031, Fig 3B, Model 5) in the clonal generations after hatching. However, we did see a significant difference between clonal isolates (df=1, *χ*^2^ =16.79, p=4.166e-05, Model 5). When tested alone, D8515 lineages that emerged early have fewer embryos compared to late hatchers (Fig 3B). However, there are relatively fewer D8515 replicates than D8222, which could be driving the clonal differences in the transgenerational models.

### Buoyancy as a possible alternative

*Daphnia* ephippia can either float to the surface or remain submerged in the water column. One concern of this experiment was that exposure to the air on the surface would mimic desiccation, causing embryos to resume development after being submerged in the 96 well plates. We found that while there was a significant effect of buoyancy on the timing of diapause termination, individuals that were fully submerged when collected from the mesocosms were more likely to emerge early than expected by chance (df=1, *χ*^2^ =8.556, p=0.003445, Model 6, Fig 4). This effect appeared to be driven by D8515, where few individuals that were floating hatched early (N=8/107). When testing D8222 alone, there was no significant difference in early diapause termination between individuals found floating or sinking (df=1, *χ*^2^ =1.528, p=0.2165). Thus, ephippia floating on the surface did not appear to drive the early termination of diapause we observed.

**Figure 4:**
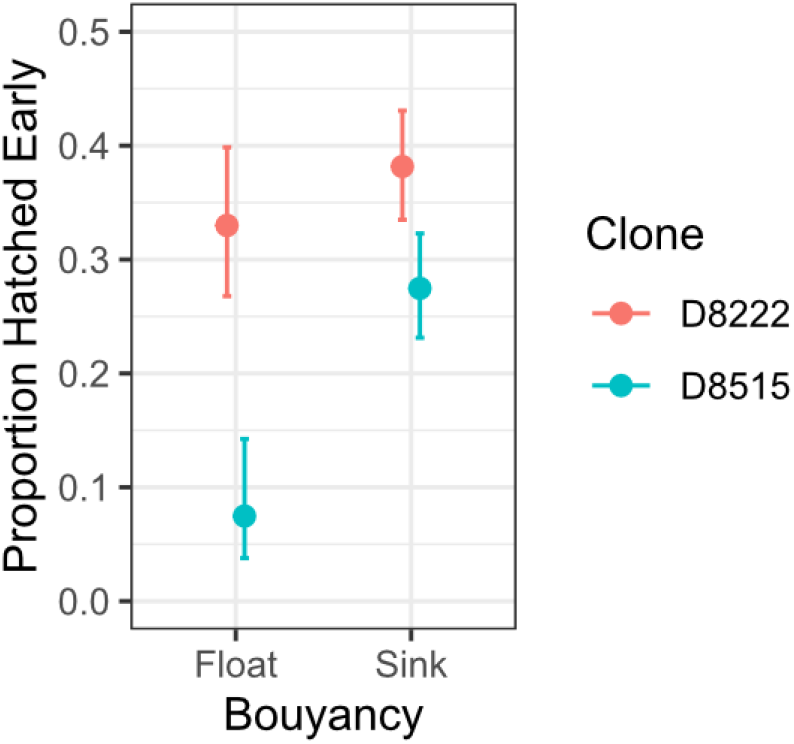
Floating ephippia does not account for early hatching. The rate of early hatching differed between epphipia found floating or sinking (df=1, *χ*^2^ =8.556, p=0.003445, Model 6). Contrary to our initial hypothesis, epphipia that were found sinking had a higher rate of early diapause termination then those found floating.

## Discussion

In this study, we examined variation in the timing of dormancy, and assessed the effects of early hatching on life history traits. We found that variation in the timing of dormancy termination in *D. pulex* is driven by maternal effects. This study contributes to the existing literature of maternally derived variation in diapause duration and provides insight into how early diapause termination may drive some transgenerational effects on reproduction.

### Early diapause termination

Our primary result is that a large proportion of hatchlings (N=413/1561, 26.46%) of a lineage of *D. pulex* collected from a semi-permanent pond emerged from diapause without any obvious changes in environmental cues. Variation in dormancy strategy has been observed in many species living in temporally heterogeneous environments, such that individuals from the same population terminate dormancy at different times (Donohue, 2009; Martínez-Ruiz & García-Roger, 2015; Polačik et al., 2017).

Variation in the duration of diapause has often been viewed as a bet-hedging strategy in both plants and animals (Evans and Dennehey 2005, Simmons 2011, Gremmer and Venable 2014). Bet-hedging is a risk-management strategy that reduces variance in mean fitness, to increase fitness over time (Simons, 2011).

Environmental variation may select for either conservative or distributive bet-hedging (Botero et al., 2015). In conservative bet-hedging, individual organisms sacrifice some immediate fitness to hedge against overall fitness loss. For example, flowering phenology could occur at an earlier point in time to hedge against an erratic end to the growing season even if it results in a lower seed set (Simons & Johnston, 2003). In distributive bet-hedging, risk management takes the form of investment into multiple strategies: for example, a female may produce offspring with a wide range of phenotypes to prepare offspring for multiple possible environments (Evans & Dennehy, 2005). Likewise, organisms of a single lineage may vary in the timing of dormancy termination to increase the chances that some offspring emerge at the right time (Sasaki & Ellner, 1995). Multiple strategies can coexist within a population, and can be driven by either genetic (Alonso-Blanco et al., 2003) or maternal effects (Martínez-Ruiz & García-Roger, 2015).

We propose that early hatching in *D. pulex* is a method for organisms to spread the risk of unsuccessful hatching, due to the unpredictability of their environment (Vanoverbeke and De Meester 2009). Due to the semi-permanent nature of our study pond, it would be advantageous for some *Daphnia* to emerge spontaneously, either as a response to better make use of a mild off-season or because they may not receive the correct cue to terminate dormancy. If individuals do not hatch out for a period of time, there is also the risk of being buried in sediment, which drastically decreases the rate of ephippial hatching (Radzikowski et al., 2016). As individuals produced are also sexually recombinant, early hatchers may add necessary genetic diversity to a mostly clonal population late in the season in permanent ponds. The increased rate of sexual reproduction in these clonal populations may provide novel gene combinations that can adapt to changes in the environment, such as increasing parasitism (Hamilton et al., 1990) and predation (Koch et al., 2020).

### Maternal control of early diapause termination

Our data suggests that there is strong maternal control of diapause variation (Fig. 1A), shown by the observation that siblings from the same ephippium tended to emerge around the same time, even after being separated. This result is consistent with prior studies linking embryonic dormancy to changes in the maternal environment. In plants such as *Campanula americana*, germination time can vary as a function of the parental environments (Galloway, 2001). In invertebrates, maternal effects have been demonstrated to influence offspring’s traits associated with dormancy. Our results resemble a previous study in a rotifer population on the effects of the maternal environment, -and subsequent maternal control, on the variation in diapause timing. (Martínez-Ruiz & García-Roger 2015). In these rotifers, the length of diapause was correlated with maternal age, and eggs produced by younger mothers had a higher chance of being late hatchers that required environmental cues to break diapause. Our results demonstrate a similar phenomenon in *Daphnia*. Individuals produced in the early stages of the mesocosm experiment, when mothers are younger, had a lower rate of early hatching (Fig. 1B).

Although we demonstrate that diapause termination is likely under maternal control, we cannot rule out genetic variation as an additional source of variation. Lineages were maintained under laboratory conditions for two years prior to starting the mesocosm cultures and mesocosms were initiated with multiple females from each clonal lineage (Barnard-Kubow et al., 2022). This might have been ample time for novel mutations to appear prior to the start of the mesocosm or during the multigeneration mesocosm experiment itself (Flynn et al., 2017; Keith et al., 2016). In principle, these newly arisen mutations could affect diapause duration and therefore be misinterpreted as maternal effects. We suggest that this scenario is unlikely for several reasons. First, it would suggest that the mutational target size for diapause termination is large. Although many genes have been implicated in diapause in invertebrates (Schwarzenberger, Chen, & Weiss, 2020), it would be difficult to imagine that it is so easy to genetically alter diapause duration. Second, most new mutations are recessive and if a mother carrying a single copy of a new mutation affecting diapause duration mated with a wild-type male, none of her offspring would manifest the phenotype. Therefore, we consider the model of genetic variation within the clonal mesocosm affecting diapause termination rate unlikely.

### Life history traits

Our analysis showed that early hatching had no significant effect on offspring survival (Fig. 2A). However, there were observed impacts on reproduction (Fig. 2B). Individuals that emerged early took slightly longer to reach reproductive age than individuals that emerged late. Coupled with the reduction in the number of resting eggs produced in subsequent generations (Fig. 3A), we hypothesize that individuals that emerged early may focus more on growth than reproduction.

For organisms that emerge from dormancy at different times, the environment that they emerge into may favor alternative life history strategies. As ephippia are typically produced late in the growing season (Altermatt & Ebert, 2008), when population density is high, then any resting eggs that emerge early would likely experience these same conditions. To survive harsh conditions, individuals that emerge into an end-of-season environment would likely be favored if they are more focused on growth/survival, rather than rapid reproduction (Mueller & Ayala, 1981). Alternatively, organisms that terminate dormancy in response to strict environmental signals, emerging later, likely experience an environment that is more hospitable with a low density of intraspecifics (Cáceres, 1998). Under low intraspecific competition, organisms that reproduce rapidly would have an advantage (Pianka, 1970). Similar trade-offs associated with size and time to reproduction have been documented as being both genetically (Reznick, 1983) and maternally (Arnold et al., 2018) derived.

## Conclusion

Our results demonstrate that *D. pulex* ephippia can hatch without experiencing changing environmental cues. The timing of emergence from dormancy also appears to be maternally derived, with siblings from the same ephippia more often hatching at the same time even after separation. We observe transgenerational effects on reproduction dependent on when individuals emerged from dormancy, with individuals that emerged early delaying reproduction. We hypothesize that early hatching may be a priming effect to survive against high intraspecific competition later in the growing season. Further research must be conducted in multiple lineages and populations to determine whether early hatching affects population dynamics in ponds.

## Statements and Declarations

### Data Availability

The datasets generated during and/or analyzed during the current study are available in the BerglandLab-Mesocosm-2019 repository, https://github.com/rjp5nc/BerglandLab-Mesocosm-2019.

## References

Alonso-Blanco, C., Bentsink, L., Hanhart, C. J., Vries, H. B., & Koornneef, M. (2003). Analysis of Natural Allelic Variation at Seed Dormancy Loci of Arabidopsis thaliana. Genetics, 164(2), 711–729. https://doi.org/10.1093/genetics/164.2.711

Altermatt, F., Ebert, D. The influence of pool volume and summer desiccation on the production of the resting and dispersal stage in a Daphnia metapopulation. Oecologia 157, 441–452 (2008). https://doi.org/10.1007/s00442-008-1080-4

Arnold, L. M., Smith, W. D., Spencer, P. D., Evans, A. N., Heppell, S. A., & Heppell, S. S. (2018). The role of maternal age and context-dependent maternal effects in the offspring provisioning of a long-lived marine teleost. Royal Society Open Science, 5(1), 170966. https://doi.org/10.1098/rsos.170966

Barnard-Kubow, K. B., Becker, D., Murray, C. S., Porter, R., Gutierrez, G., Erickson, P., Nunez, J. C. B., Voss, E., Suryamohan, K., Ratan, A., Beckerman, A., & Bergland, A. O. (2022). Genetic Variation in Reproductive Investment Across an Ephemerality Gradient in Daphnia pulex. Molecular Biology and Evolution, 39(6), msac121. https://doi.org/10.1093/molbev/msac121

Botero, C. A., Weissing, F. J., Wright, J., & Rubenstein, D. R. (2015). Evolutionary tipping points in the capacity to adapt to environmental change. Proceedings of the National Academy of Sciences, 112(1), 184–189. https://doi.org/10.1073/pnas.1408589111

ten Brink, H., Gremer, J. R., & Kokko, H. (2020). Optimal germination timing in unpredictable environments: The importance of dormancy for both among- and within-season variation. Ecology Letters, 23(4), 620–630. https://doi.org/10.1111/ele.13461

Cáceres, C. E. (1998). Interspecific Variation in the Abundance, Production, and Emergence of Daphnia Diapausing Eggs. Ecology, 79(5), 1699–1710. https://doi.org/10.1890/0012-9658(1998)079[1699:IVITAP]2.0.CO;2

Davison, J. (1969). Activation of the Ephippial Egg of Daphnia pulex. The Journal of General Physiology, 53(5), 562–575.

Dolfi, L., Ripa, R., Antebi, A., Valenzano, D. R., & Cellerino, A. (2019). Cell cycle dynamics during diapause entry and exit in an annual killifish revealed by FUCCI technology. EvoDevo, 10(1), 29. https://doi.org/10.1186/s13227-019-0142-5

Donohue, K. (2009). Completing the cycle: Maternal effects as the missing link in plant life histories. Philosophical Transactions of the Royal Society B: Biological Sciences, 364(1520), 1059–1074. https://doi.org/10.1098/rstb.2008.0291

Ensslin, A., Van de Vyver, A., Vanderborght, T., & Godefroid, S. (2018). Ex situ cultivation entails high risk of seed dormancy loss on short-lived wild plant species. Journal of Applied Ecology, 55(3), 1145–1154. https://doi.org/10.1111/1365-2664.13057

Evans, M. E. K., & Dennehy, J. J. (2005). GERM BANKING: BET-HEDGING AND VARIABLE RELEASE FROM EGG AND SEED DORMANCY. The Quarterly Review of Biology, 80(4), 431–451. https://doi.org/10.1086/498282

Flynn, J. M., Chain, F. J. J., Schoen, D. J., & Cristescu, M. E. (2017). Spontaneous Mutation Accumulation in Daphnia pulex in Selection-Free vs. Competitive Environments. Molecular Biology and Evolution, 34(1), 160–173. https://doi.org/10.1093/molbev/msw234

Galloway, L. F. (2001). The effect of maternal and paternal environments on seed characters in the herbaceous plant Campanula Americana (Campanulaceae). American Journal of Botany, 88(5), 832–840. https://doi.org/10.2307/2657035

Gerber, N., Kokko, H., Ebert, D., & Booksmythe, I. (2018). Daphnia invest in sexual reproduction when its relative costs are reduced. Proceedings of the Royal Society B: Biological Sciences, 285(1871), 20172176. https://doi.org/10.1098/rspb.2017.2176

Gliwicz, Z. M., & Lampert, W. (1994). Clutch-size variability in Daphnia: Body-size related effects of egg predation by cyclopoid copepods. Limnology and Oceanography, 39(3), 479–485. https://doi.org/10.4319/lo.1994.39.3.0479

Grosbois, G., Mariash, H., Schneider, T., & Rautio, M. (2017). Under-ice availability of phytoplankton lipids is key to freshwater zooplankton winter survival. Scientific Reports, 7(1), Article 1. https://doi.org/10.1038/s41598-017-10956-0

Hairston, N. G., & Kearns, C. M. (2002). Temporal Dispersal: Ecological and Evolutionary Aspects of Zooplankton Egg Banks and the Role of Sediment Mixing. Integrative and Comparative Biology, 42(3), 481–491.

Hamilton, W. D., AXELRODtt, R., & Tanese, R. (1990). Sexual reproduction as an adaptation to resist parasites (a review). Proc. Natl. Acad. Sci. USA, 8.

Hand, S. C., Denlinger, D. L., Podrabsky, J. E., & Roy, R. (2016). Mechanisms of animal diapause: Recent developments from nematodes, crustaceans, insects, and fish. American Journal of Physiology - Regulatory, Integrative and Comparative Physiology, 310(11), R1193–R1211. https://doi.org/10.1152/ajpregu.00250.2015

Hebert, P. D. N., & Ward, R. D. (1972). Inheritance during Parthenogenesis in DAPHNIA MAGNA. Genetics, 71(4), 639–642.

Hiruta, C., & Tochinai, S. (2014). Formation and structure of the ephippium (resting egg case) in relation to molting and egg laying in the water flea Daphnia pulex De Geer (Cladocera: Daphniidae). Journal of Morphology, 275(7), 760–767. https://doi.org/10.1002/jmor.20255

Keith, N., Tucker, A. E., Jackson, C. E., Sung, W., Lledó, J. I. L., Schrider, D. R., Schaack, S., Dudycha, J. L., Ackerman, M., Younge, A. J., Shaw, J. R., & Lynch, M. (2016). High mutational rates of large-scale duplication and deletion in Daphnia pulex. Genome Research, 26(1), 60–69. https://doi.org/10.1101/gr.191338.115

Koch, H. R., Wagner, S., & Becks, L. (2020). Antagonistic species interaction drives selection for sex in a predator– prey system. Journal of Evolutionary Biology, 33(9), 1180–1191. https://doi.org/10.1111/jeb.13658

Lopes, F. L., Desmarais, J. A., & Murphy, B. D. (2004). Embryonic diapause and its regulation. Reproduction, 128(6), 669–678. https://doi.org/10.1530/rep.1.00444

Luu, D. H.-K., Vo, H. P., & Xu, S. (2020). An Efficient Method for Hatching Diapausing Embryos of Daphnia pulex Species Complex (Crustacea, Anomopoda). Journal of Experimental Zoology. Part A, Ecological and Integrative Physiology, 333(2), 111–117. https://doi.org/10.1002/jez.2331

Lynch, M. (1983). Ecological Genetics of Daphnia Pulex. Evolution, 37(2), 358–374. https://doi.org/10.1111/j.1558-5646.1983.tb05545.x

Martínez-Ruiz, C., & García-Roger, E. M. (2015). Being first increases the probability of long diapause in rotifer resting eggs. Hydrobiologia, 745(1), 111–121. https://doi.org/10.1007/s10750-014-2098-8

Mueller, L. D., & Ayala, F. J. (1981). Trade-off between r-selection and K-selection in Drosophila populations. Proceedings of the National Academy of Sciences of the United States of America, 78(2), 1303–1305. https://doi.org/10.1073/pnas.78.2.1303

Paes, T. A. S. V., Rietzler, A. C., Pujoni, D. G. F., & Maia-Barbosa, P. M. (2016). High temperatures and absence of light affect the hatching of resting eggs of Daphnia in the tropics. Anais Da Academia Brasileira De Ciencias, 88(1), 179–186. https://doi.org/10.1590/0001-3765201620140595

Pancella, J. R., & Stross, R. G. (1963). Light induced hatching ofDaphnia resting eggs. Chesapeake Science, 4(3), 135–140. https://doi.org/10.2307/1350746

Pianka, E. R. (1970). On r- and K-Selection. The American Naturalist, 104(940), 592–597.

Polačik, M., Smith, C., & Reichard, M. (2017). Maternal source of variability in the embryo development of an annual killifish. Journal of Evolutionary Biology, 30(4), 738–749. https://doi.org/10.1111/jeb.13038

Postma, F. M., & Ågren, J. (2022). Effects of primary seed dormancy on lifetime fitness of Arabidopsis thaliana in the field. Annals of Botany, 129(7), 795–808. https://doi.org/10.1093/aob/mcac010

Radzikowski, J., Krupińska, K., & Ślusarczyk, M. (2018). Different thermal stimuli initiate hatching of Daphnia diapausing eggs originating from lakes and temporary waters. Limnology, 19(1), 81–88. https://doi.org/10.1007/s10201-017-0520-4

Radzikowski, J., Sikora, A., & Ślusarczyk, M. (2016). The effect of lake sediment on the hatching success of Daphnia ephippial eggs. Journal of Limnology, 75(3), Article 3. https://doi.org/10.4081/jlimnol.2016.1345

Reznick, D. (1983). The Structure of Guppy Life Histories: The Tradeoff between Growth and Reproduction. Ecology, 64(4), 862–873. https://doi.org/10.2307/1937209

Ricci, C. (2001). Dormancy patterns in rotifers. Hydrobiologia, 446(1), 1–11. https://doi.org/10.1023/A:1017548418201

Sasaki, A., & Ellner, S. (1995). The Evolutionarily Stable Phenotype Distribution in a Random Environment. Evolution, 49(2), 337–350. https://doi.org/10.1111/j.1558-5646.1995.tb02246.x

Satake, A., Sasaki, A., & Iwasa, Y. (2001). Variable timing of reproduction in unpredictable environments: Adaption of flood plain plants. Theoret. Popul. Biol, 1–15.

Schwartz, S. S., & Hebert, P. D. N. (1987). Methods for the activation of the resting eggs of Daphnia. Freshwater Biology, 17(2), 373–379. https://doi.org/10.1111/j.1365-2427.1987.tb01057.x

Schwarzenberger, A., Chen, L. & Weiss, L.C. The expression of circadian clock genes in Daphnia magna diapause. Sci Rep 10, 19928 (2020). https://doi.org/10.1038/s41598-020-77065-3

Shabani, A., Shabanpour, B., & Hoseini, S. (2012). Hatching Requirements of Daphnia magna Straus, 1820, and Daphnia pulex Linnaeus, 1758, Diapausing Eggs from Iranian Populations In vitro. Journal of Agricultural Science and Technology, 14.

Simons, A. M. (2011). Modes of response to environmental change and the elusive empirical evidence for bet hedging. Proceedings of the Royal Society B: Biological Sciences, 278(1712), 1601–1609. https://doi.org/10.1098/rspb.2011.0176

Simons, A. M., & Johnston, M. O. (2003). Suboptimal timing of reproduction in Lobelia inflata may be a conservative bet-hedging strategy. Journal of Evolutionary Biology, 16(2), 233–243. https://doi.org/10.1046/j.1420-9101.2003.00530.x

Standard, A. (2007) Standard guide for conducting acute toxicity tests on test materials with fishes, macroinvertebrates, and amphibians. West Conshohocken, PA, U. S. DOI 01520E0729-96

Vandekerkhove, J., Declerck, S., Brendonck, L., Conde-Porcuna, J. M., Jeppesen, E., Johansson, L. S., & De Meester, L. (2005). Uncovering hidden species: Hatching diapausing eggs for the analysis of cladoceran species richness. Limnology and Oceanography: Methods, 3(9), 399–407. https://doi.org/10.4319/lom.2005.3.399

Wagner, I., & Simons, A. M. (2009). Divergence in germination traits among arctic and alpine populations of Koenigia islandica: Light requirements. Plant Ecology, 204(1), 145–153. https://doi.org/10.1007/s11258-009-9578-3

Winsor, G. L., & Innes, D. J. (2002). Sexual reproduction in Daphnia pulex (Crustacea: Cladocera): observations on male mating behaviour and avoidance of inbreeding. Freshwater Biology, 47(3), 441–450. https://doi.org/10.1046/j.1365-2427.2002.00817.x

